# RHO-1 and the Rho GEF RHGF-1 interact with UNC-6/Netrin signaling to regulate growth cone protrusion and microtubule organization in *C. elegans*

**DOI:** 10.1101/520262

**Authors:** Mahekta R. Gujar, Aubrie M. Stricker, Erik A. Lundquist

## Abstract

UNC-6/Netrin is a conserved axon guidance cue that directs growth cone migrations in the dorsal-ventral axis of *C. elegans* and in the vertebrate spinal cord. UNC-6/Netrin is expressed in ventral cells, and growth cones migrate ventrally toward or dorsally away from UNC-6/Netrin. Recent studies of growth cone behavior during outgrowth *in vivo* in *C. elegans* have led to a polarity/protrusion model in directed growth cone migration away from UNC-6/Netrin. In this model, UNC-6/Netrin first polarizes the growth cone via the UNC-5 receptor, leading to dorsally biased protrusion and F-actin accumulation. UNC-6/Netrin then regulates protrusion based on this polarity. The receptor UNC-40/DCC drives protrusion dorsally, away from the UNC-6/Netrin source, and the UNC-5 receptor inhibits protrusion ventrally, near the UNC-6/Netrin source, resulting in dorsal migration. UNC-5 inhibits protrusion in part by excluding microtubules from the growth cone, which are pro-protrusive. Here we report that the RHO-1/RhoA GTPase and its activator GEF RHGF-1 inhibit growth cone protrusion and MT accumulation in growth cones, similar to UNC-5. However, growth cone polarity of protrusion and F-actin were unaffected by RHO-1 and RHGF-1. Thus, RHO-1 signaling acts specifically as a negative regulator of protrusion and MT accumulation, and not polarity. Genetic interactions suggest that RHO-1 and RHGF-1 act with UNC-5, as well as with a parallel pathway, to regulate protrusion. The cytoskeletal interacting molecule UNC-33/CRMP was required for RHO-1 activity to inhibit MT accumulation, suggesting that UNC-33/CRMP might act downstream of RHO-1. In sum, these studies describe a new role of RHO-1 and RHGF-1 in regulation of growth cone protrusion by UNC-6/Netrin.

**Author Summary:** Neural circuits are formed by precise connections between axons. During axon formation, the growth cone leads the axon to its proper target in a process called axon guidance. Growth cone outgrowth involves asymmetric protrusion driven by extracellular cues that stimulate and inhibit protrusion. How guidance cues regulate growth cone protrusion in neural circuit formation is incompletely understood. This work shows that the signaling molecule RHO-1 acts downstream of the UNC-6/Netrin guidance cue to inhibit growth cone protrusion in part by excluding microtubules from the growth cone, which are structural elements that drive protrusion.

## Introduction

The connectivity of neuronal circuits is established through properly guided axons which form functional synaptic connections. The growing axon is guided to its target by the motile, actin-based growth cone at the tip of the growing neurite. Growth cone response to extracellular guidance cues allows the axon to extend, retract, turn and branch, regulated by the reorganization and dynamics of the actin and microtubule cytoskeletons of the growth cone (Dent and Gertler 2003).

In *C. elegans* and vertebrates, the conserved laminin-like UNC-6/Netrin guidance cue and its receptors UNC-40/DCC and UNC-5 direct dorsal-ventral axon outgrowth (Hedgecock *et al.* 1990; Ishii *et al.* 1992; Leung-Hagesteijn *et al.* 1992; Chan *et al.* 1996; Leonardo *et al.* 1997; Hong *et al.* 1999; Montell 1999; Shekarabi and Kennedy 2002; Moore *et al.* 2007). UNC-6 is secreted by cells in the ventral nerve cord (Wadsworth *et al.* 1996), and growth cones grow toward UNC-6/Netrin (i.e. ventral migration; attraction) and away from UNC-6/Netrin (i.e. dorsal migration; repulsion). The prevailing model of UNC-6/Netrin-mediated axon guidance involves a ventral-to-dorsal chemotactic gradient of the molecule, which growth cones interpret by migrating up or down the gradient using the “attractive” receptor UNC-40/DCC or the “repulsive” receptor UNC-5, respectively (Tessier-Lavigne and Goodman 1996; Lai Wing Sun *et al.* 2011). However, this model has recently been challenged by studies in mouse spinal cord showing that floorplate Netrin is dispensable for commissural axon guidance, and that ventricular expression is important, possibly in a close-range, haptotactic event (Dominici *et al.* 2017; Varadarajan and Butler 2017; Varadarajan *et al.* 2017; Yamauchi *et al.* 2017).

Recent studies of growth cones during their formation and outgrowth *in vivo* in *C. elegans* suggest that UNC-40/DCC and UNC-5 each act in both attracted and repelled growth cones. In the HSN neuron, which extends an axon ventrally, UNC-6/Netrin controls the ventral accumulation of the UNC-40 receptor in the HSN cell body, and UNC-5 acts to focus UNC-40/DCC ventrally (the statistically-oriented asymmetric localization (SOAL) model) (Kulkarni *et al.* 2013; Yang *et al.* 2014; Limerick *et al.* 2017). Furthermore, our previous work with the VD growth cones that migrate dorsally (repelled) suggests that UNC-6/Netrin first polarizes protrusion and F-actin to the dorsal side of the growth cone via the UNC-5 receptor, and then regulates protrusion based on this polarity (the polarity/protrusion model). UNC-5 inhibits protrusion ventrally, close to the UNC-6/Netrin source, and UNC-40 stimulates protrusion dorsally, away from the UNC-6/Netrin source, resulting in directed dorsal growth away from UNC-6/Netrin (Norris and Lundquist 2011; Norris *et al.* 2014; Gujar *et al.* 2018).

UNC-40/DCC drives growth cone lamellipodial and filopodial protrusion via the small GTPases CDC-42, CED-10/Rac, and MIG-2/RhoG, the Rac-specific guanine nucleotide exchange factor (GEF) TIAM-1, and actin cytoskeletal regulators Arp2/3, UNC-34/Enabled and UNC-115/abLIM (Gitai *et al.* 2003; Struckhoff and Lundquist 2003; Shakir *et al.* 2008; Norris *et al.* 2009; Demarco *et al.* 2012). UNC-5 inhibits growth cone protrusion via the Rac GEF UNC-73/trio, CED-10/Rac and MIG-2/RhoG (also used to drive protrusion), the FMO flavin monooxygenases which might act via actin, and the actin and MT-interacting proteins UNC-33/CRMP and UNC-44/Ankyrin (Norris and Lundquist 2011; Norris *et al.* 2014; Gujar *et al.* 2017). UNC-5 also restricts the accumulation of microtubule + ends in VD growth cones which have pro-protrusive effects (Gujar *et al.* 2018). Thus, in *unc-5* mutants, VD growth cones are larger and more protrusive, display unpolarized protrusion including ventral protrusions, display unpolarized F-actin around the periphery of the growth cone, and have increased accumulation of MT + ends (Norris and Lundquist 2011; Gujar *et al.* 2018). This unregulated protrusion results in unfocused growth cones that fail to migrate dorsally away from UNC-6/Netrin, causing the severe VD axon guidance defects seen in *unc-5* mutants.

The Rho-family GTPases CED-10/Rac, MIG-2/RhoG, and CDC-42 control neuronal protrusion (Lundquist *et al.* 2001; Struckhoff and Lundquist 2003; Demarco *et al.* 2012; Norris *et al.* 2014). Here we dissect the role of RHO-1, the single RhoA molecule encoded in the *C. elegans* genome, in regulation of VD growth cone polarity and protrusion. *rho-1* RNAi results in early embryonic arrest, with a failure in cytokinesis and severe morphological defects (Spencer *et al.* 2001; Bringmann and Hyman 2005; Morita *et al.* 2005; Motegi and Sugimoto 2006). We used cell-specific expression of constitutively-active RHO-1(G14V) and dominant-negative RHO-1(T19N), and cell-specific RNAi of *rho-1* and found that RHO-1 inhibited growth cone protrusion and MT + end accumulation. RHO-1 did not, however, affect polarity of protrusion or F-actin. We also found that the RHO-1 activator RHGF-1, a RHO-1 GTP exchange factor of the LARG family (Yau *et al.* 2003; Chen *et al.* 2014), was required to inhibit protrusion and MT + end accumulation similar to RHO-1. Genetic interactions with UNC-5 signaling and UNC-33/CRMP suggest that RHGF-1 and RHO-1 might act downstream of UNC-5 and in parallel to other regulators of protrusion and MT + end accumulation. These studies also revealed that RHO-1 requires UNC-33/CRMP to prevent MT + end accumulation. In sum, results reported here show that RHGF-1 and RHO-1 are key inhibitors of growth cone protrusion and MT + end accumulation and act with UNC-5 in protrusion, but not growth cone polarity.

## Results

### RHO-1 regulates growth cone protrusion but not polarity

RHO-1 is the single RhoA homolog in *C. elegans.* Loss of *rho-1* leads to embryonic lethality, with a failure in cytokinesis (Jantsch-Plunger *et al.* 2000), and perturbation of RHO-1 signaling in adults results in dysfunction in numerous neuronal and non-neuronal functions leading to death (McMullan and Nurrish 2011). To understand the role of RHO-1 in VD growth cone morphology, we constructed constitutively-active G14V and dominant-negative T19N versions of RHO-1, and expressed them in the VD/DD neurons using the *unc-25* promoter. Constitutively-active *rho-1(G14V)* expression significantly reduced the VD growth cone area and shortened filopodial protrusions as compared to wild-type (Figure 1A-B, D). In contrast, dominant-negative *rho-1(T19N)* expression displayed significantly longer filopodial protrusions as compared to wild-type VD growth cones (Figure 1A-B, E). Growth cone area was increased, but not significantly so. These results indicate that RHO-1 activity inhibits growth cone protrusion.

**Figure 1.**
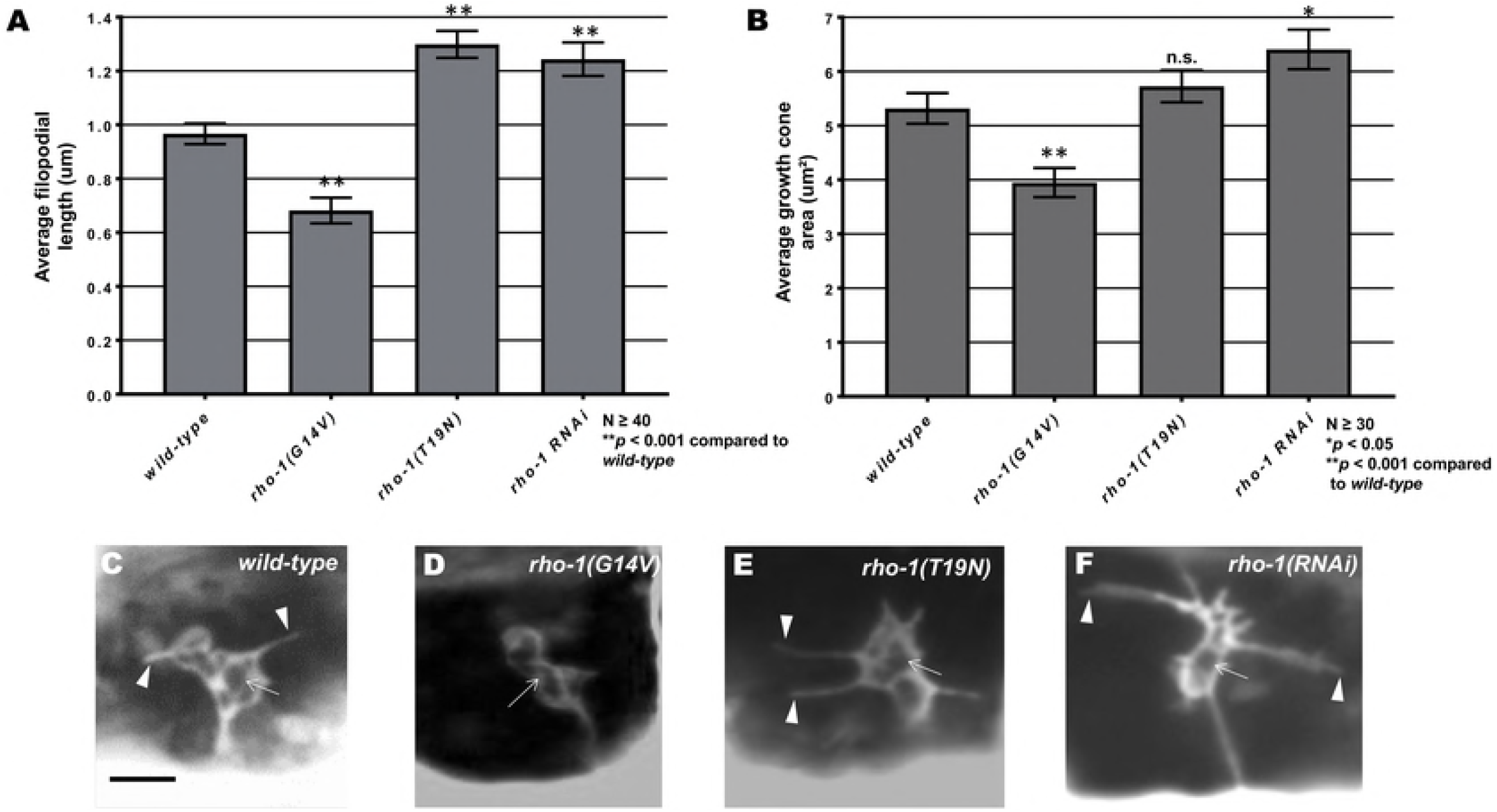
VD growth cone protrusion and polarity in *rho-1* mutants. (A-B) Quantification of VD growth cone filopodial length and growth cone area in wild-type and *rho-1* mutant animals (See Materials and Methods). (A) Average filopodial length, in μm. (B) Growth cone area in μm^2^. Error bars represent 2x standard error of the mean; asterisks indicate the significant difference between wild-type and the mutant phenotype (\**p* < 0.05, \*\**p* < 0.001) determined by two-sided t-test with unequal variance. n.s., not significant. (C-E) Fluorescence micrographs of VD growth cones with *Punc-25::gfp* expression *(juIs76*); (C) A wild-type VD growth cone. (D) *rho-1(G14V)* showing small and inhibited VD growth cone phenotype (E) *rho-1(T19N)* and (F) *rho-1(RNAi)* growth cones showing increased filopodial protrusion in the form of longer filopodia. Arrows point to the growth cone and arrow heads indicate representative filopodia. (G) A graph showing the percent of dorsally-directed filopodial protrusions in VD growth cones of different genotypes (see Materials and Methods). (H-I) VD growth cones with *Punc-25::gfp* expression *(juIs76*). The solid horizontal lines indicate the dorsal and ventral extent of the growth cone body, and the hatched lines indicate the average center of the growth cone. Protrusions above the hatched horizontal line are considered dorsal, and those below ventral. Scale bars represent 5μm.

We used a transgenic RNAi approach to knock down *rho-1* in the VD/DD motor neurons as previously described (see Materials and Methods) (Esposito *et al.* 2007; Sundararajan *et al.* 2014). Plasmids were generated to drive expression of sense and antisense RNA fragments complementary to the *rho-1* under the control of the *unc-25* promoter. Animals were made transgenic with a mix of the sense and antisense plasmids, and the resulting transgenes were used in analysis. The average length of filopodial protrusions and growth cone area were significantly increased in *rho-1(RNAi)* (Figure 1A-B, F). These data suggest that RHO-1 normally inhibits VD growth cone protrusion.

The polarity of filopodial protrusions was not affected by *rho-1(DN)* or *rho-1(RNAi),* as protrusions still displayed a dorsal bias similar wild-type (Figure 1G-I). Thus, despite showing increased protrusion, the polarity of growth cone protrusion was not affected by *rho-1.*

### RHO-1 is required to limit EBP-2::GFP puncta accumulation in VD growth cones

Previous studies indicate that in VD growth cones, F-actin accumulates at the dorsal, protrusive edge of the growth cone and acts as a polarity mark to specify protrusion in this region (Figure 2A and B) (Norris and Lundquist 2011; Gujar *et al.* 2018). Furthermore, microtubule + ends are present in the growth cone and have a pro-protrusive role (Gujar *et al.* 2018). In wild-type, MT + ends are rare in VD growth cones (~2 per growth cone) (Figure 2E and F) (Gujar *et al.* 2018), and protrusion is tightly regulated and localized to the dorsal leading edge at the site of F-actin accumulation Figure 2) (Gujar *et al.* 2018).

VD growth cone F-actin was monitored using the VAB-1ABD::GFP reporter, and MT + ends were monitored using EBP-2::GFP as described previously (Norris and Lundquist 2011; Gujar *et al.* 2018). Dominant-negative *rho-1(T19N)* and *rho-1(RNAi)* had no effect on dorsally-polarized F-actin accumulation (Figure 2A and D), consistent with no effects on growth cone polarity of protrusion (Figure 1). However, growth cone EBP-2::GFP puncta number were significantly increased by dominant-negative *rho-1(T19N)* and *rho-1(RNAi)* (Figure 2E, G, and H), consistent with increased protrusion in these backgrounds.

Constitutively-active *rho-1(G14V)* resulted in fewer EBP-2::GFP puncta, consistent with reduced growth cone protrusion (Figure 2E). F-actin polarity was also abolished, with distribution along the periphery of the entire growth cone (Figure 2A and C). Possibly, constitutive activation reveals a role of RHO-1 in F-actin polarity that is not affected in reduction of function treatments. However, a similar effect on F-actin was observed with constitutively-active Rac GTPases MIG-2 and CED-10 (Gujar *et al.* 2018). Possibly, this effect on F-actin is a consequence of small growth cones with severely-restricted protrusion, and not a direct role in F-actin organization. In sum, these results suggest that RHO-1 normally restricts growth cone protrusion by preventing accumulation of growth cone MT + ends.

### The RhoGEF RHGF-1 acts with RHO-1 to inhibit growth cone filopodial protrusion and MT + end accumulation

RHGF-1 is a PDZ RhoGEF with PDZ, RGS, C1, DH, and PH domains (Figure 3A). RHGF-1 is a RHO-1-specific GEF and acts with RHO-1 in neurotransmitter release and axonal regeneration (Yau *et al.* 2003; Hiley *et al.* 2006; Lin *et al.* 2012; Chen *et al.* 2014; Alam *et al.* 2016). *rhgf-1(ok880)* is a 1170bp in frame deletion which removes a large part of the DH domain and is predicted to have no RhoGEF activity (Hiley et al., 2006), *rhgf-1(gk217)* is a 247bp in frame deletion which removes the C1 domain, and *rhgf-1(gk292502)* produces a premature stop just before the C1 domain (Figure 3A). *rhgf-1* mutants each displayed increased growth cone area and longer filopodial protrusions compared to wild-type (Figure 3B-F). The dorsally-biased polarity of growth cone protrusion was not significantly affected by *rhgf-1* mutation (Figure 3G-I). These data indicate that RHGF-1 is normally required to limit the extent of growth cone protrusion, but does not regulate growth cone polarity, similar to *rho-1.*

The *Drosophila* RHGF-1 homolog DRhoGEF2 is a key regulator of morphogenesis and associates with the tips of growing MTs and exhibits plus end tracking (Rogers *et al.* 2004). In *C. elegans,* RHGF-1 associates with MTs and initiates an axon regeneration pathway (Chen et al., 2014). *rhgf-1* mutant VD growth cones displayed significantly increased numbers of EBP-2::GFP puncta (Figure 4A-C), but caused no significant defects in F-actin organization, similar to *rho-1* knockdown (Figure 4D-F). These results indicate that RHGF-1 might act with RHO-1 to inhibit growth cone protrusion by excluding MT+ ends from entering the growth cone periphery.

The results above indicate that the VD growth cones of activated *rho-1(G14V)* displayed reduced protrusion, and that those of *rhgf-1* loss of function were overly-protrusive. The VD growth cones of activated *rho-1(G14V)* double mutants with *rhgf-1* loss of function resembled the small, inhibited growth cones of *rho-1(G14V)* alone (Figure 5A-E), with a significant reduction in filopodial length and growth cone area as compared to *wild-type* and *rhgf-1* mutants alone (Figure 5A-E). Similarly, double mutants of *rhgf-1* and *rho-1(G14V)* showed a significant decrease in the average number of EBP-2 puncta in the growth cone similar to *rho-1(G14V)* alone (Figure 6A-D). VAB-10ABD::GFP distribution in these double mutant growth cones also resembled activated *rho-1(G14V)* with F-actin distributed randomly all across the growth cone (Figure 6E-H). That activated RHO-1(G14V) was epistatic to *rhgf-1* loss of function is consistent with RHO-1 acting downstream of RHGF-1 in limiting growth protrusion and EBP-2 accumulation in VD growth cones.

### Activated *myr::unc-40* and *myr::unc-5* require RHGF-1

Previous studies showed that UNC-6/Netrin signaling via the heterodimeric UNC-40/UNC-5 receptor is required for inhibition of growth cone protrusion in UNC-6/Netrin repulsive axon guidance (Norris and Lundquist 2011; Norris *et al.* 2014). Constitutive activation of UNC-40 and UNC-5 using myristoylated versions of the cytoplasmic domains of UNC-40 and UNC-5 *(myr::unc-40* and *myr::unc-5)* in the VD neurons result in small growth cones with few or no filopodial protrusions (Gitai *et al.* 2003; Norris and Lundquist 2011; Norris *et al.* 2014). Loss of *rhgf-1* significantly suppressed inhibition of filopodial protrusion and growth cone size caused by *myr::unc-40* and *myr::unc-5* (Figure 7).

*myr::unc-40* and *myr::unc-5* growth cones show a significant decrease in the average number of EBP-2::GFP puncta in the VD growth cones as compared to wild-type (Figure 8A-C) (Gujar *et al.* 2018). Double mutants of *rhgf-1* with *myr::unc-40* and *myr::unc-5* resembled *rhgf-1* mutants alone, with significant increases in protrusion and MT+ end accumulation (Figure 8A, D). Similar to activated Racs and RHO-1(G14V), F-actin is distributed throughout the small growth cones in activated *myr::unc-5* and *myr::unc-40* (Figure 8E-G). *rhgf-1* mutation restored dorsal polarity of F-actin (Figure 8E and H). In sum, the growth cones of *rhgf-1* double mutants with *myr::unc-5* and *myr::unc-40* resembled those of *rhgf-1* mutants alone: increased protrusion and EBP-2 puncta accumulation, but normal dorsal F-actin polarity. These data indicate that RHGF-1 is required for the inhibitory effects of *myr::unc-40* and *myr::unc-5* on growth cone protrusion and EBP-2::GFP puncta accumulation.

### Activated RHO-1 does not suppress *unc-5* loss of function

*unc-5* loss of function results in unpolarized, overly-protrusive VD growth cones. Excess MT + ends accumulate in *unc-5,* and dorsal polarity of F-actin accumulation and thus protrusion is lost (Norris and Lundquist 2011; Gujar *et al.* 2018). Activated *rho-1(G14V)* expression did not suppress the large growth cone area and long filopodial protrusions seen in *unc-5* mutants (i.e. double mutants resembled *unc-5* alone) (Figure 9). Furthermore, we observed no significant change in EBP-2::GFP and VAB-10ABD::GFP distribution in the VD growth cones as compared to *unc-5* mutants alone (Figures 10 and 11).

### Activated RHO-1 suppresses *unc-33/CRMP* loss of function

The Collapsin-response mediator protein (CRMP) UNC-33 and the Ankyrin-like molecule UNC-44 are required for inhibition of growth cone protrusion of activated *myr::unc-40* and *myr::unc-5.* Loss of *unc-33* and *unc-44* results in VD growth cones resembling *unc-5* mutants, with increased protrusion, increased MT + end accumulation, and loss of F-actin dorsal polarity (Norris *et al.* 2014; Gujar *et al.* 2018).

Double mutants of *unc-33* and *rho-1(G14V)* resembled those of activated *rho-1(G14V)* mutants alone, with a significant decrease in growth cone area and filopodial protrusions (Figure 11). Despite reduced protrusion and smaller growth cone size, EBP-2::GFP puncta accumulation was increased in double mutants of *unc-33* and *rho-1(G14V)* (Figure 12). By contrast, double mutants of *unc-44* with *rho-1(G14V)* resembled *unc-44* mutants, with excessive growth cone filopodial as evidenced with increased filopodial length and growth cone area, as well as an increase in EBP-2 puncta distribution (Figures 11 and 12). Double mutants of *unc-33* and *unc-44* with *rho-1(G14V)* showed no significant change in F-actin distribution as compared to single mutants alone (Figure 13). These complex interactions reveal a differentiation of function between UNC-33/CRMP and UNC-44/Ankyrin in interaction with RHO-1 in growth cone morphology regulation.

## Discussion

Previous studies indicate that directed outgrowth of the VD growth cones away from UNC-6/Netrin involves a polarity/protrusion mechanism (Norris and Lundquist 2011; Norris *et al.* 2014; Gujar *et al.* 2018). UNC-6/Netrin first polarizes protrusion to the dorsal side of the growth cone, and then regulates the extent of growth cone protrusion, with the receptor UNC-40 stimulating protrusion dorsally and the UNC-5 receptor inhibiting protrusion ventrally, resulting in directed dorsal growth away from UNC-6/Netrin. Growth cone polarity is reflected in F-actin polarity, with F-actin distribution biased to the dorsal side of the growth cone (i.e. the protrusive side). Growth cone protrusion correlates with the presence of MT + ends, and MTs are pro-protrusive in the VD growth cones (Gujar *et al.* 2018). UNC-6/Netrin, its receptors UNC-5 and UNC-40, Rac GTPases, and UNC-33/CRMP all regulate both growth cone polarity and protrusion (Norris and Lundquist 2011; Norris *et al.* 2014; Gujar *et al.* 2018). UNC-5 and UNC-33 normally inhibit growth cone protrusion in part by restricting MT + end accumulation in growth cones (Gujar *et al.* 2018).

Our results here show that the small GTPases RHO-1 and the Rho Guanine nucleotide Exchange factor RHGF-1 mediate inhibition of growth cone protrusion and are required to limit MT+ -end accumulation in growth cones, similar to UNC-5 and UNC-33. However, RHO-1 and RHGF-1 had no effect on growth cone polarity (i.e. mutants did not affect dorsally-biased distribution of filopodial protrusion and F-actin). Thus, RHO-1 and RHGF-1 specifically affect VD growth cone protrusion, and not polarity. Activated RHO-1 was epistatic to *rhgf-1* loss of function (i.e. growth cones in double mutants displayed inhibited filopodial protrusions and a significant reduction in EBP-2 puncta distribution similar to activated *rho-1* alone), consistent with the known role of RHGF-1 as an upstream Rho activator.

Genetic studies suggest a complex interaction of RHO-1 and RHGF-1 with UNC-5 and UNC-33. The data are consistent with the idea that RHO-1 and RHGF-1 act in the UNC-5 pathway as well as in a parallel pathway (RHGF-1 was required for the effects of activated MYR::UNC-5, but activated RHO-1 did not suppress *unc-5* loss of function) (Figure 13). Additionally, activated RHO-1 suppressed the large, protrusive growth cones of *unc-33* loss-of-function, but did not decrease MT + end accumulation in these small growth cones. This suggests that UNC-33 might act downstream of RHO-1 in MT accumulation, and that RHO-1 has an UNC-33-independent role in protrusion. While we do not fully understand the nature of these interactions at this point, our data clearly show that RHO-1 and RHGF-1 interact with UNC-6/Netrin signaling to regulate growth cone protrusion and MT organization during growth cone outgrowth.

### RHO-1 regulates growth cone protrusion and EBP-2 distribution

Expression of activated RHO-1(G14V) resulted in VD growth cones with a marked decrease in growth cone protrusion and EBP-2 puncta distribution (Figures 1 and 2). Expression of the dominant negative form of RHO-1(T19N) in the VD neurons and *rho-1(RNAi)* resulted in increased protrusion and EBP-2::GFP accumulation. MT+ – ends in the growth cone periphery (Figures 1 and 2). Notably, neither dominant-negative RHO-1(T19N) or *rho-1(RNAi)* resulted in altered growth cone polarity and F-actin dorsal bias (Figures 1 and 2), suggesting that RHO-1 might specifically affect growth cone protrusion but not polarity.

Previous work has identified roles of the Rho GTPases in regulation of both microtubules and actin (Wittmann and Waterman-Storer 2001). RhoA has been shown to regulate formation of contractile actin structures such as stress fibers and promote stabilization of microtubules (Cook *et al.* 1998; Etienne-Manneville and Hall 2002) through actomyosin contraction. In cultured growth cones, RhoA is involved in F-actin retrograde flow, wherein actin filaments in the periphery undergo constant retrograde transport to growth cone body (Lin and Forscher 1995; Lin *et al.* 1996; Zhang *et al.* 2003; Van Goor *et al.* 2012). RhoA activates RhoA kinase (ROCK), which activates contractility by phosphorylating the regulatory myosin light chain (MLC). This actin retrograde flow is thought to restrict MTs from the growth cone through physical association with these actin filaments undergoing retrograde flow, thereby reducing leading edge protrusion resulting in growth cone collapse and retraction (ZHANG *et al.* 2003; Gallo 2004). Growth cone advance can occur when this actin-MT linkage is disrupted or when actin becomes attached to the substrate (the “clutch” hypothesis) (Nichol *et al.* 2016) resulting in anterograde flow over the anchored actin filaments. One hypothesis explaining our results is that, in VD growth cones, RHO-1-mediated retrograde flow of actin restricts MT + ends from the growth cones, and when RHO-1 activity is reduced, more MTs enter the growth cones resulting in increased growth cone protrusion.

### The Rho GEF RHGF-1 acts with RHO-1 to inhibit growth cone protrusion and MT accumulation

Loss of *rhgf-1* resulted in increased growth cone protrusion and accumulation of EBP-2::GFP, similar to but more pronounced than dominant-negative RHO-1(T19N) and *rho-1(RNAi)* (Figures 3 and 4). Furthermore, *rhgf-1* mutants had no effect on growth cone polarity of protrusion or F-actin distribution (Figure 4). RHGF-1 might be an activator of RHO-1 to inhibit growth cone protrusion and MT accumulation. Consistent with this idea, activated *rho-1* was epistatic to *rhgf-1* loss-of-function (i.e. activating RHO-1 bypasses the need for RHGF-1). Growth cones in these double mutants displayed inhibited protrusion and reduction in MT distribution similar to activated *rho-1* alone, suggesting that RHGF-1 acts as an upstream RHO-1 regulator in this process (Figure 5 and 6).

Previous studies in *Drosophila* s2 cells have shown that the RHGF-1 homolog, DRhoGEF2, induces contractile cell shape changes by regulating myosin II dynamics via Rho1 pathway. Furthermore, DRhoGEF2 associates with tips of growing MTs and travels to the cell cortex (Rogers *et al.* 2004). In *C. elegans,* RHGF-1 functions through Rho and ROCK to activate the MAPKKK DLK-1 during MT disruption, triggering synaptic branch retraction and overgrowth of PLM neurites ultimately leading to neuronal remodeling (Chen *et al.* 2014). Possibly, RHGF-1 activates RHO-1 to mediate a potential retrograde flow of F-actin to restrict MT accumulation in the growth cone.

### RHGF-1 is required for the inhibitory effects of MYR::UNC-5 and MYR::UNC-40

*rhgf-1* loss-of-function suppressed the inhibitory effects of activated *myr::unc-40* and *myr::unc-5* on growth cones. Double mutant growth cones resembled those of *rhgf-1* alone, with increased protrusion and EBP-2::GFP puncta (Figure 7 and 8). That RHGF-1 is required for the effects of constitutively active MYR::UNC-40 and MYR::UNC-5 suggest that RHGF-1 acts downstream of MYR::UNC-5 and MYR::UNC-40. However, it is possible that RHGF-1 defines a parallel pathway. In any event, the inhibitory effects of MYR::UNC-5 and MYR::UNC-40 require functional RHGF-1.

### Activated RHO-1(G14V) cannot compensate for loss of UNC-5 in growth cone inhibition

Receptors to several attractive or repulsive guidance cues signal through complex pathways through the Rho family of small GTPases to direct changes in growth cone cytoskeletal organization (Luo 2002; Govek *et al.* 2005), and Rho activity is thought to be induced by “repulsive” cues (Guan and Rao 2003). Loss of the UNC-6/Netrin receptor *unc-5* has been shown to cause excessively large VD growth cones with increased protrusion and excess EBP-2::GFP accumulation (Norris and Lundquist 2011; Gujar *et al.* 2018).

If RHO-1 is activated by UNC-5, we expect that activated *rho-1(G14V)* would be epistatic to *unc-5* loss-of-function. This was not the case, as growth cones of *rho-1(G14V); unc-5(lof)* double mutants resembled those of *unc-5(lof)* alone, with increased protrusiveness and EBP-2::GFP accumulation (Figure 9 and 10). Possibly, loss of UNC-5 affects multiple parallel pathways, including RHO-1, and activation of the RHO-1 pathway alone cannot compensate for loss of UNC-5. Alternately, RHO-1 might act in parallel to UNC-5. That RHGF-1 function is required for the effects of activated MYR::UNC-5 and MYR::UNC-40 suggests that RHGF-1 (and by extension RHO-1) might, in part, act in the UNC-5 pathway directly.

### UNC-33/CRMP is required for activated RHO-1(G14V) restriction of EBP-2::GFP

Previous studies have shown that the *C. elegans* UNC-33/CRMP is required in a pathway downstream with Rac GTPases for inhibition of growth cone protrusion in response to UNC-6/Netrin (Norris *et al.* 2014). *unc-33* loss-of-function mutants show large protrusive growth cones with excess EBP-2 accumulation in the growth cones, similar to *unc-5.* While activated RHO-1(G14V) did not suppress the excessively-protrusive growth cones of *unc-5* mutants, it did suppress those of *unc-33* (Figure 11). Protrusion of growth cones of *rho-1(G14V); unc-33* double mutants resembled *rho-1(G14V)* alone (i.e. protrusion was reduced and growth cones were small).

Interestingly, despite their small size, inhibited *unc-33; rho-1(G14V)* growth cones displayed increased EBP-2 puncta compared to wild-type animals, but significantly lower than *unc-33* mutants alone (Figure 12). Thus, activated RHO-1(G14V) can fully suppress excess protrusion, but not EBP-2::GFP accumulation, of *unc-33* mutants. Together, these results suggest that UNC-33 is required for activated RHO-1(G14V) to restrict MTs from growth cones. They also suggest that RHO-1 has a role in protrusion that is independent of MT accumulation, as protrusion was reduced in *rho-1(G14V); unc-33* double mutants despite excess MT accumulation.

UNC-44/Ankyrin is required to properly localize UNC-33/CRMP to the axons (Maniar *et al.* 2011), and mutants are phenotypically indistinguishable in the VD growth cones (both are required to polarize protrusion and F-actin and to inhibit protrusion and EBP-2::GFP accumulation) (Norris *et al.* 2014; Gujar *et al.* 2018). However, *unc-44* loss was completely epistatic to activated RHO-1(G14V), including both protrusion and EBP-2::GFP accumulation. This suggests that UNC-44/Ankyrin has a role that is independent of UNC-33/CRMP involving non-MT-based regulation of protrusion. The FMO flavin monooxygenases inhibit growth cone protrusion with UNC-5 (Gujar *et al.* 2017), possibly in an actin-based manner similar to MICAL (Hung *et al.* 2010; Hung *et al.* 2011). Possibly, UNC-44/Ankyrin acts in this pathway or another independently from UNC-33/CRMP.

## Summary

Our results show that RHO-1 and the Rho activator GEF RHGF-1 are required to inhibit VD growth cone protrusion and to restrict EBP-2::GFP puncta accumulation in growth cones, possibly downstream of the UNC-6/Netrin receptor UNC-5. One potential scenario for how these molecules interact is shown in Figure 13. UNC-5 might activate RHGF-1 and thus RHO-1, and UNC-33/CRMP might then be required to exclude MTs from growth cones in response to RHO-1 activation.

CRMP interactions with Rho, actin, and microtubules have been documented in other systems. In cultured mammalian neurons, CRMP interacts with F-actin and with tubulin dimers to promote microtubule assembly (Fukata *et al.* 2002; Rosslenbroich *et al.* 2005), and expression of CRMP2 can alter Rho-GTPase-driven neurite morphology. Co-expression of Crmp-2 with activated Rho can promote cell spreading and neurite growth and this function of Crmp-2 is regulated by Rho Kinase (Hall *et al.* 2001). Furthermore, CRMP-2 has been shown to be phosphorylated by Rho Kinase II (Arimura *et al.* 2000; Arimura *et al.* 2005) which disrupts the association of mature full-length CRMP-2 with tubulin heterodimers so that tubulin cannot be transported to the plus ends of microtubules for assembly (Fukata *et al.* 2002) causing neurite retraction and growth cone collapse (Arimura and Kaibuchi 2007). This reduced binding capacity to tubulin by phosphorylated CRMP-2, can be reversed by inhibiting RhoA activity (Petratos *et al.* 2008). Thus, RHO-1 may regulate growth cone protrusion and MT distribution through the phosphorylation activity of UNC-33/CRMP possibly through the same pathway or in parallel to it.

If RHO-1 is indeed involved in F-actin retrograde flow, the role of UNC-33 might be to link F-actin to microtubules, such that in an *unc-33* mutant, MTs are not excluded despite retrograde flow (including in the activated RHO-1(G14V) background). RHO-1 might have an additional non-UNC-33 and non-MT-dependent role in inhibiting protrusion, along with UNC-44, possibly involving actin. In sum, RHO-1 is a key negative regulator of growth cone protrusion and MT accumulation that acts specifically in the protrusion aspect of the polarity/protrusion model of directed growth cone migration away from UNC-6/Netrin.

## Materials and methods

### Genetic methods

Experiments were performed at 20°C using standard *C. elegans* techniques (Brenner 1974). Mutations used were LGIV: *unc-5(e53* and *e152), unc-33(e204), unc-44(e362*); *lqls128 [Punc-25::myr::unc-40]* LGX: *rhgf-1(gk217, ok880 and gk292502), lqls170 [rgef-1::vab-10ABD::gfp].* Chromosomal locations not determined: *lqIs279 [Punc-25::ebp-2::gfp]* by integration of *lqEx809, lhIs6 [Punc-25::mCherry], lqIs296 [Punc-25::myr::unc-5], lqIs312 [Punc-25::rho-1(G14V)]* by integration of *lqEx1043, lqIs314 [Punc-25::rho-1(T19N)]* by integration of *lqEx1070.* Extrachromosomal arrays were generated using standard gonadal injection (Mello and Fire, 1995) and include: *lqEx999* and *lqEx1000 [Punc-25::myr::unc-40; Pgcy-32::yfp], lqEx1131, lqEx1132, lqEx1133* and *lqEx1134 [Punc-25::rho-1 RNAi; Pgcy-32::yfp], OX347 [Prgef-1::vab-10ABD::gfp; ttx-3::rfp].* Multiple (≥3) extrachromosomal transgenic lines of *Punc-25::ebp-2::gfp, Punc-25::rho-1(G14V)* and *Punc-25::rho-1(T19N)* were analyzed with similar effect, and one was chosen for integration and further analysis.

### Growth cone imaging

VD growth cones were imaged and quantified as previously described (Norris and Lundquist 2011). Briefly, animals at ~16 h post-hatching at 20°C were placed on a 2% agarose pad and paralyzed with 5mM sodium azide in M9 buffer, which was allowed to evaporate for 4 min before placing a coverslip over the sample. Some genotypes were slower to develop than others, so the 16 h time point was adjusted for each genotype. Growth cones were imaged with a Qimaging Rolera mGi camera on a Leica DM5500 microscope. Images were analyzed in ImageJ, and statistical analyses done with Graphpad Prism software. As described in (Norris and Lundquist 2011; Norris *et al.* 2014), growth cone area was determined by tracing the perimeter of the growth cone body, not including filopodia. Average filopodial length was determined using a line tool to trace the length of the filopodium. Unless otherwise indicated, ≥25 growth cones were analyzed for each genotype. These data were gathered in ImageJ and entered into Graphpad Prism for analysis. A two-sided t-test with unequal variance was used to determine significance of difference between genotypes.

### VAB-10ABD::GFP imaging

The F-actin binding domain of VAB-10/spectraplakin fused to GFP has been used to monitor F-actin in *C. elegans* (Bosher *et al.* 2003; Patel *et al.* 2008). We used it to image F-actin in the VD growth cones as previously described (Norris and Lundquist 2011). To control for variability in growth cone size and shape, and as a reference for asymmetric localization of VAB-10ABD::GFP, a soluble mCherry volume marker was included in the strain. Growth cones images were captured as described above. ImageJ was used image analysis to determine asymmetric VAB-10ABD::GFP localization. For each growth cone, five line scans were made from dorsal to ventral. For each line, pixel intensity was plotted as a function of distance from the dorsal leading edge of the growth cone. The average intensity (arbitrary units) and standard error for each growth cone was determined. For dorsal versus ventral comparisons, the pixel intensities for VAB-10ABD::GFP were normalized to the volumetric mCherry fluorescence in line scans from the dorsal half and the ventral half of each growth cone. This normalized ratio was determined for multiple growth cones, and the average and standard error for multiple growth cones was determined. Statistical comparisons between genotypes were done using a two-tailed t-test with unequal variance on these average normalized ratios of multiple growth cones of each genotype.

### EBP-2::GFP imaging

EBP-2::GFP has previously been used to monitor microtubule plus ends in other *C. elegans* cells including neurons (Srayko *et al.* 2005; Kozlowski *et al.* 2007; Yan *et al.* 2013). We constructed a transgene consisting of the *unc-25* promoter driving expression of *ebp-2::gfp* in the VD/DD neurons. In growth cones, a faint fluorescence was observed throughout the growth cone, resembling a soluble GFP and allowing for the growth cone perimeter to be defined. In addition to this faint, uniform fluorescence, brighter puncta of EBP-2::GFP were observed that resembled the EBP-1::GFP puncta described in other cells and neurons. For each growth cone, the perimeter and filopodia were defined, and the EBP-2::GFP puncta in the growth cone were counted. For each genotype, the puncta number for many growth cones (≥25 unless otherwise noted) was determined. Puncta number displayed high variability within and between genotypes, so box-and-whiskers plots (Graphpad Prism) were used to accurately depict this variation. The grey boxes represent the upper and lower quartiles of the data set, and the “whiskers” represent the high and low values. Dots represent major outliers. Significance of difference was determined by a two-sided t-test with unequal variance.

### Transgenic RNA-mediated gene interference (RNAi)

We used a cell-specific transgenic RNAi approach as described previously (Esposito *et al.* 2007). Fragments of the *rho-1* coding region was amplified by PCR and inserted behind the *unc-25* promoter in a plasmid (primer and plasmid sequences available upon request). A “sense” and “antisense” orientation relative to the *unc-25* promoter was isolated. An equimolar mixture of the sense and antisense plasmids was used to construct transgenic animals. These transgenic animals were predicted to express both sense and antisense RNAs driven by the *unc-25* promoter in the VD/DD motor neurons, which was expected to trigger a double-stranded RNA response in these cells (RNAi).

## Acknowledgments

The authors thank the members of the Lundquist and Ackley labs for discussion and E. Struckhoff for technical assistance. Some strains were provided by the CGC, which is funded by NIH Office of Research Infrastructure Programs (P40 0D010440).

This work was supported by NIH grants R01NS040945, R56NS095682 and P20GM103638. A.M.S. was a Kansas Infrastructure Network of Biomedical Excellence Undergraduate Scholar and Star Trainee (NIH P20GM103418) and was supported by the University of Kansas Center for Undergraduate Research.

---

**Figure 2. VD growth cone F-actin polarity and EBP-2::GFP accumulation in *rho-1* mutants.** (A) The average dorsal/ventral ratio of GFP/mCherry from multiple growth cones in wild-type and mutant animals expressing VAB-10ABD::GFP and mCherry (a volumetric marker) as described previously (Norris and Lundquist 2011) (see Materials and Methods) Error bars represent 2x standard error of the mean. Asterisks (*) indicate the significant difference between wild-type and the mutant phenotype (\**p* < 0. 05) determined by two-sided t-test with unequal variance. (B-D) Representative images of VD growth cones with cytoplasmic mCherry in red (a volumetric marker) and VAB-10ABD::GFP in green. Areas of overlap are yellow (arrows). Dashed lines indicate the growth cone periphery. Dorsal is up and anterior is left. Scale bar: 5 μm. (B) A wild-type VD growth cone, (C) *rho-1(G14V)* showing an inhibited growth cone with F-actin accumulation all along the growth cone and (D) *rho-1(T19N)* VD growth cones with VAB-10ABD::GFP expression in the dorsal leading edge of the growth cone. (E) Box-and-whiskers plot of the number of EBP-2::GFP puncta in the growth cones of different genotypes (≥25 growth cones for each genotype). The grey boxes represent the upper and lower quartiles, and error bars represent the upper and lower extreme values. Dots represent outliers. Asterisks (*) indicate the significant difference between wild-type and the mutant phenotype (\*\**p* < 0.001) determined by two-sided t-test with unequal variance. n.s., not significant. (F-H) Fluorescence micrographs of EBP-2 distribution in the VD growth cones; (F) A wild-type VD growth cone and (G) *rho-1(T19N)* and (H) *rho-1(RNAi)* growth cones showing increased puncta in the growth cone and filopodial protrusions. Arrows indicate representative EBP-2::GFP puncta. Dashed lines indicate the growth cone perimeter. Dorsal is up and anterior is left. Scale bar: 5μm.

**Figure 3. Growth cone protrusion and polarity in *rhgf-1* loss-of-function.** (A) A schematic diagram of the predicted 1,340-amino acid residue RHGF-1 molecule. PDZ= PDZ domain, RGS= Regulator of G protein signaling domain, C1= Ester/diacylglycerol binding domain, DH= Dbl homology domain, PH= Plekstrin homology domain. Extent of deletions of *ok880* and *gk217* are indicated the red lines. The red arrow points to the premature stop site in *gk292502.* (B-C) Quantification of VD growth cone filopodial length and growth cone area as described in Figure 1. \**p* < 0.05 and \*\**p* < 0.001, determined by two-sided t-test with unequal variance. n.s., not significant. (D-F) Fluorescence micrographs of VD growth cones *(juIs76[Punc-25::gfp]).* Arrows point to the growth cone and arrow heads indicate representative filopodia. Scale bar: 5μm. (G) A graph showing the percent of dorsally-directed filopodial protrusions in VD growth cones of different genotypes as described in Figure 1. (H-I) Growth cone polarity of protrusion as described in Figure 1.

**Figure 4. EBP-2::GFP accumulation and F-actin polarization in *rhgf-1* mutants.** (A) Quantification of the number of EBP-2::GFP puncta in wild-type and *rhgf-1* mutant growth cones as described in Figure 2E. Asterisks (*) indicate the significant difference between wild-type and the mutant phenotype (\**p* < 0.05,\*\**p* < 0.001) determined by twosided t-test with unequal variance. n.s., not significant. (B-C) Fluorescence micrographs of EBP-2 distribution in the VD growth cones; (B) A wild-type VD growth cone (C) *rhgf-1(gk292502)* growth cones showing increased puncta in the growth cone and filopodial protrusions. Arrows indicate representative EBP-2::GFP puncta. Dashed lines indicate the growth cone perimeter. Dorsal is up and anterior is left. Scale bar: 5μm. (D) The average dorsal-to-ventral ratio of VAB-10ABD::GFP/mCherry from multiple growth cones in wild-type and mutant animals as described in Figure 2A. Error bars represent 2x standard error of the mean; n.s., not significant. (E-F) Representative images of VD growth cones with cytoplasmic mCherry in red (a volumetric marker) and the VAB-10ABD::GFP in green. Areas of overlap are yellow (arrows). Dashed lines indicate the growth cone periphery. Dorsal is up and anterior is left. Scale bar: 5 μm. (E) A wild-type growth cone and (F) *rhgf-1(ok880)* growth cones with VAB-10ABD::GFP expression in the dorsal leading edge of the growth cone.

**Figure 5. Genetic interactions of *rhgf-1* and *rho-1* in growth cone protrusion.** (A-B) Quantification of VD growth cone filopodial length and growth cone area in single and double mutant animals as described in Figure 1. (A) Average filopodial length, in μm. (B) Growth cone area in μm^2^. Error bars represent 2x standard error of the mean; asterisks indicate the significant difference between *rhgf-1* single mutants and the double mutant phenotype (\*\**p* < 0.001) determined by two-sided t-test with unequal variance. (C-E) Fluorescence micrographs of VD growth cones as described in Figure 1. Arrows point to the growth cone and arrow heads indicate representative filopodia. Scale bar: 5μm.

**Figure 6. Genetic interactions of *rhgf-1* and *rho-1* in EBP-2::GFP accumulation and F-actin polarization. (A)** Quantification of the number of EBP-2::GFP puncta in wild-type and mutant animals as described in Figure 2E. Asterisks (*) indicate the significant difference between *rhgf-1* single mutants and the double mutant phenotype (\*\**p* < 0.01) determined by two-sided t-test with unequal variance. (B-D) Fluorescence micrographs of EBP-2 distribution in the VD growth cones; (B) *rhgf-1(ok880)* growth cone (C) *rho-1(G14V)* and (D) *rhgf-1(ok880); rho-1(G14V)* growth cones with decreased *ebp-2* puncta. Arrows indicate representative EBP-2::GFP puncta. Dashed lines indicate the growth cone periphery. Dorsal is up and anterior is left. Scale bar: 5 μm. (E) The average dorsal-to-ventral ratio of GFP/mCherry from multiple growth cones in single and double mutant animals as described in Figure 2A. (F-H) Representative images of VD growth cones with cytoplasmic mCherry in red (a volumetric marker) and the VAB-10ABD::GFP in green. Areas of overlap are yellow (arrows). Error bars represent 2x standard error of the mean. Asterisks (*) indicate the significant difference between single and double mutant phenotype (\**p* < 0.05) determined by two-sided t-test with unequal variance. Dashed lines indicate the growth cone periphery. Dorsal is up and anterior is left. Scale bar: 5 μm.

**Figure 7. Genetic interactions of *rhgf-1* with *myr::unc-40* and *myr::unc-5* in growth cone protrusion.** (A) Quantification of VD growth cone filopodial length and growth cone area in single and double mutant animals as described in Figure 1. (A) Average filopodial length, in μm. (B) Growth cone area in μm^2^. Error bars represent 2x standard error of the mean; asterisks indicate the significant difference between *myr::unc-40,* single and double mutants (\*\**p* < 0.001) determined by two-sided t-test with unequal variance. (C-E) Fluorescence micrographs of mutant VD growth cones as described in Figure 1. Arrows point to the growth cone and arrow heads indicate representative filopodia. Scale bar: 5μm.

**Figure 8. Genetic interactions of *rhgf-1* with *myr::unc-40* and *myr::unc-5* in EBP-2::GFP accumulation and F-actin polarity.** (A) Quantification of the number of EBP-2::GFP puncta in wild-type and mutant animals as described in Figure 2E. Asterisks (*) indicate the significant difference between *myr::unc-40,* single mutants and double mutants (\*\**p* < 0.001), determined by two-sided t-test with unequal variance. (B-E) Fluorescence micrographs of EBP-2 distribution in the VD growth cones. Arrows indicate representative EBP-2::GFP puncta. Dashed lines indicate the growth cone periphery. Dorsal is up and anterior is left. Scale bar: 5 μm. (E) The average dorsal-to-ventral ratio of GFP/mCherry from multiple growth cones in wild-type, single and double mutant animals as described in Figure 2A. Error bars represent 2x standard error of the mean. Asterisks (*) indicate the significant difference between *myr::unc-40,* single mutants and double mutants (\**p* < 0.05) determined by two-sided t-test with unequal variance. (F-H) Representative images of VD growth cones with cytoplasmic mCherry in red (a volumetric marker) and the VAB-10ABD::GFP in green. Areas of overlap are yellow (arrows). Dashed lines indicate the growth cone periphery. Dorsal is up and anterior is left. Scale bar: 5μm.

**Figure 9. Genetic interactions of *rho-1* and *unc-5* in growth cone protrusion.** (A-B) Quantification of VD growth cone filopodial length and growth cone area in single and double mutant animals as described in Figure 1. (A) Average filopodial length, in μm. (B) Growth cone area in μm^2^. Error bars represent 2x standard error of the mean; n.s., not significant determined by two-sided t-test with unequal variance. (C-E) Fluorescence micrographs of mutant VD growth cones. Arrows point to the growth cone and arrow heads indicate representative filopodia. Scale bar: 5μm.

**Figure 10. Genetic interactions of *rho-1* and *unc-5* in EBP-2::GFP accumulation and F-actin polarity.** (A) Quantification of the number of EBP-2::GFP puncta in wild-type and mutant animals as described in Figure 2E. n.s., not significant, determined by two-sided t-test with unequal variance. (B-D) Fluorescence micrographs of EBP-2 distribution in the VD growth cones. Arrows indicate representative EBP-2::GFP puncta. Dashed lines indicate the growth cone periphery. Dorsal is up and anterior is left. Scale bar: 5 μm. (E) The average dorsal-to-ventral ratio of GFP/mCherry from multiple growth cones in wild-type, single and double mutant animals as described in Figure 2A. Error bars represent 2x standard error of the mean; n.s. indicates no significant difference between *unc-5* single mutants and double mutants determined by two-sided t-test with unequal variance. (F-H) Representative images of VD growth cones with cytoplasmic mCherry in red (a volumetric marker) and the VAB-10ABD::GFP in green. Areas of overlap are yellow (arrows). (F) wild-type growth cone, (G) *rho-1(G14V)* growth cone, (H) *unc-5(e53); rho-1(G14V)* double mutant VD growth cones with cytoplasmic mCherry and VAB-10ABD::GFP expression. Dashed lines indicate the growth cone periphery. Dorsal is up and anterior is left. Scale bar: 5 μm.

**Figure 11. Genetic interaction *rho-1* with *unc-33* and *unc-44* in growth cone protrusion.** (A-B) Quantification of VD growth cone filopodial length and growth cone area in single and double mutant animals as described in Figure 1. (A) Average filopodial length, in μm. (B) Growth cone area in μm^2^. Error bars represent 2x standard error of the mean; asterisks indicate the significant difference between the single mutant and the double mutant phenotype (\*\**p* < 0.001) determined by two-sided t-test with unequal variance. n.s., not significant. (C-E) Fluorescence micrographs of mutant VD growth cones as described in Figure 1. Arrows point to the growth cone and arrow heads indicate representative filopodia. Scale bar: 5μm.

**Figure 12. Genetic interactions of RHO-1 and *unc-33* and *unc-44* in EBP-2::GFP accumulation and F-actin polarity.** (A) Quantification of the number of EBP-2::GFP puncta in wild-type and mutant animals as described in Figure 2E. Asterisks (*) indicate the significant difference between single mutants and the double mutant (\**p* < 0.01). Pound (#) indicates significant difference between wild-type and double mutant (#*p* < 0.01) determined by two-sided t-test with unequal variance. (B-D) Fluorescence micrographs of EBP-2 distribution in the VD growth cones. (B) An *unc-33(e204)* growth cone with increased *ebp-2* puncta. (D) A *rho-1(G14V)* small and inhibited growth cone with significantly fewer *ebp-2* puncta. (E) An *unc-33(e204); rho-1(G14V)* small and inhibited growth cone with increased *ebp-2* puncta. Arrows indicate representative EBP-2::GFP puncta. Dashed lines indicate the growth cone periphery. Dorsal is up and anterior is left. Scale bar: 5 μm. (E) The average dorsal-to-ventral ratio of GFP/mCherry from multiple growth cones in wild-type as described in Figure 2E. Error bars represent 2x standard error of the mean; n.s. indicates no significant difference between *unc-33* and *unc-44* single mutants and their respective double mutants determined by twosided t-test with unequal variance. (F-H) Representative images of VD growth cones with cytoplasmic mCherry in red (a volumetric marker) and the VAB-10ABD::GFP in green. Areas of overlap are yellow (arrows). Scale bar: 5 μm. Dashed lines indicate the growth cone periphery. Dorsal is up and anterior is left.

**Figure 13. Possible interactions of RHO-1 in growth cone MT exclusion.** UNC-5 might activate RHGF-1 and thus RHO-1, and UNC-33/CRMP is required for RHO-1 activity to exclude MTs from the growth cone. UNC-5 might activate Rac GTPases in parallel to drive MT exclusion via UNC-33/CRMP. UNC-5 might also engage a parallel pathway to drive MT exclusion.

## Reference

Alam, T., H. Maruyama, C. Li, S. I. Pastuhov, P. Nix et al., 2016 Axotomy-induced HIF-serotonin signalling axis promotes axon regeneration in C. elegans. Nat Commun 7: 10388.

Arimura, N., N. Inagaki, K. Chihara, C. Menager, N. Nakamura et al., 2000 Phosphorylation of collapsin response mediator protein-2 by Rho-kinase. Evidence for two separate signaling pathways for growth cone collapse. J Biol Chem 275: 23973–23980.

Arimura, N., and K. Kaibuchi, 2007 Neuronal polarity: from extracellular signals to intracellular mechanisms. Nat Rev Neurosci 8: 194–205.

Arimura, N., C. Menager, Y. Kawano, T. Yoshimura, S. Kawabata et al., 2005 Phosphorylation by Rho kinase regulates CRMP-2 activity in growth cones. Mol Cell Biol 25: 9973–9984.

Bosher, J. M., B. S. Hahn, R. Legouis, S. Sookhareea, R. M. Weimer et al., 2003 The Caenorhabditis elegans vab-10 spectraplakin isoforms protect the epidermis against internal and external forces. J Cell Biol 161: 757–768.

Brenner, S., 1974 The genetics of Caenorhabditis elegans. Genetics 77: 71–94.

Bringmann, H., and A. A. Hyman, 2005 A cytokinesis furrow is positioned by two consecutive signals. Nature 436: 731–734.

Chan, S. S., H. Zheng, M. W. Su, R. Wilk, M. T. Killeen et al., 1996 UNC-40, a C. elegans homolog of DCC (Deleted in Colorectal Cancer), is required in motile cells responding to UNC-6 netrin cues. Cell 87: 187–195.

Chen, C. H., A. Lee, C. P. Liao, Y. W. Liu and C. L. Pan, 2014 RHGF-1/PDZ-RhoGEF and retrograde DLK-1 signaling drive neuronal remodeling on microtubule disassembly. Proc Natl Acad Sci U S A 111: 16568–16573.

Cook, T. A., T. Nagasaki and G. G. Gundersen, 1998 Rho guanosine triphosphatase mediates the selective stabilization of microtubules induced by lysophosphatidic acid. J Cell Biol 141: 175–185.

Demarco, R. S., E. C. Struckhoff and E. A. Lundquist, 2012 The Rac GTP exchange factor TIAM-1 acts with CDC-42 and the guidance receptor UNC-40/DCC in neuronal protrusion and axon guidance. PLoS Genet 8: e1002665.

Dent, E. W., and F. B. Gertler, 2003 Cytoskeletal dynamics and transport in growth cone motility and axon guidance. Neuron 40: 209–227.

Dominici, C., J. A. Moreno-Bravo, S. R. Puiggros, Q. Rappeneau, N. Rama et al., 2017 Floor-plate-derived netrin-1 is dispensable for commissural axon guidance. Nature 545: 350–354.

Esposito, G., E. Di Schiavi, C. Bergamasco and P. Bazzicalupo, 2007 Efficient and cell specific knock-down of gene function in targeted C. elegans neurons. Gene 395: 170–176.

Etienne-Manneville, S., and A. Hall, 2002 Rho GTPases in cell biology. Nature 420: 629–635.

Fukata, Y., T. J. Itoh, T. Kimura, C. Menager, T. Nishimura et al., 2002 CRMP-2 binds to tubulin heterodimers to promote microtubule assembly. Nat Cell Biol 4: 583–591.

Gallo, G., 2004 Myosin II activity is required for severing-induced axon retraction in vitro. Exp Neurol 189: 112–121.

Gitai, Z., T. W. Yu, E. A. Lundquist, M. Tessier-Lavigne and C. I. Bargmann, 2003 The netrin receptor UNC-40/DCC stimulates axon attraction and outgrowth through enabled and, in parallel, Rac and UNC-115/AbLIM. Neuron 37: 53–65.

Govek, E. E., S. E. Newey and L. Van Aelst, 2005 The role of the Rho GTPases in neuronal development. Genes Dev 19: 1–49.

Guan, K. L., and Y. Rao, 2003 Signalling mechanisms mediating neuronal responses to guidance cues. Nat Rev Neurosci 4: 941–956.

Gujar, M. R., A. M. Stricker and E. A. Lundquist, 2017 Flavin monooxygenases regulate Caenorhabditis elegans axon guidance and growth cone protrusion with UNC-6/Netrin signaling and Rac GTPases. PLoS Genet 13: e1006998.

Gujar, M. R., L. Sundararajan, A. Stricker and E. A. Lundquist, 2018 Control of Growth Cone Polarity, Microtubule Accumulation, and Protrusion by UNC-6/Netrin and Its Receptors in Caenorhabditis elegans. Genetics 210: 235–255.

Hall, C., M. Brown, T. Jacobs, G. Ferrari, N. Cann et al., 2001 Collapsin response mediator protein switches RhoA and Rac1 morphology in N1E-115 neuroblastoma cells and is regulated by Rho kinase. J Biol Chem 276: 43482–43486.

Hedgecock, E. M., J. G. Culotti and D. H. Hall, 1990 The unc-5, unc-6, and unc-40 genes guide circumferential migrations of pioneer axons and mesodermal cells on the epidermis in C. elegans. Neuron 4: 61–85.

Hiley, E., R. McMullan and S. J. Nurrish, 2006 The Galpha12-RGS RhoGEF-RhoA signalling pathway regulates neurotransmitter release in C. elegans. EMBO J 25: 5884–5895.

Hong, K., L. Hinck, M. Nishiyama, M. M. Poo, M. Tessier-Lavigne et al., 1999 A ligand-gated association between cytoplasmic domains of UNC5 and DCC family receptors converts netrin-induced growth cone attraction to repulsion. Cell 97: 927–941.

Hung, R. J., C. W. Pak and J. R. Terman, 2011 Direct redox regulation of F-actin assembly and disassembly by Mical. Science 334: 1710–1713.

Hung, R. J., U. Yazdani, J. Yoon, H. Wu, T. Yang et al., 2010 Mical links semaphorins to F-actin disassembly. Nature 463: 823–827.

Ishii, N., W. G. Wadsworth, B. D. Stern, J. G. Culotti and E. M. Hedgecock, 1992 UNC-6, a laminin-related protein, guides cell and pioneer axon migrations in C. elegans. Neuron 9: 873–881.

Jantsch-Plunger, V., P. Gonczy, A. Romano, H. Schnabel, D. Hamill et al., 2000 CYK-4: A Rho family gtpase activating protein (GAP) required for central spindle formation and cytokinesis. J Cell Biol 149: 1391–1404.

Kozlowski, C., M. Srayko and F. Nedelec, 2007 Cortical microtubule contacts position the spindle in C. elegans embryos. Cell 129: 499–510.

Kulkarni, G., Z. Xu, A. M. Mohamed, H. Li, X. Tang et al., 2013 Experimental evidence for UNC-6 (netrin) axon guidance by stochastic fluctuations of intracellular UNC-40 (DCC) outgrowth activity. Biol Open 2: 1300–1312.

Lai Wing Sun, K., J. P. Correia and T. E. Kennedy, 2011 Netrins: versatile extracellular cues with diverse functions. Development 138: 2153–2169.

Leonardo, E. D., L. Hinck, M. Masu, K. Keino-Masu, S. L. Ackerman et al., 1997 Vertebrate homologues of C. elegans UNC-5 are candidate netrin receptors. Nature 386: 833–838.

Leung-Hagesteijn, C., A. M. Spence, B. D. Stern, Y. Zhou, M. W. Su et al., 1992 UNC-5, a transmembrane protein with immunoglobulin and thrombospondin type 1 domains, guides cell and pioneer axon migrations in C. elegans. Cell 71: 289–299.

Limerick, G., X. Tang, W. S. Lee, A. Mohamed, A. Al-Aamiri et al., 2017 A Statistically Oriented Asymmetric Localization (SOAL) Model for Neuronal Outgrowth Patterning by Caenorhabditis elegans UNC-5 (UNC5) and UNC-40 (DCC) Netrin Receptors. Genetics.

Lin, C. H., E. M. Espreafico, M. S. Mooseker and P. Forscher, 1996 Myosin drives retrograde F-actin flow in neuronal growth cones. Neuron 16: 769–782.

Lin, C. H., and P. Forscher, 1995 Growth cone advance is inversely proportional to retrograde F-actin flow. Neuron 14: 763–771.

Lin, L., T. Tran, S. Hu, T. Cramer, R. Komuniecki et al., 2012 RHGF-2 is an essential Rho-1 specific RhoGEF that binds to the multi-PDZ domain scaffold protein MPZ-1 in Caenorhabditis elegans. PLoS One 7: e31499.

Lundquist, E. A., P. W. Reddien, E. Hartwieg, H. R. Horvitz and C. I. Bargmann, 2001 Three C. elegans Rac proteins and several alternative Rac regulators control axon guidance, cell migration and apoptotic cell phagocytosis. Development 128: 4475–4488.

Luo, L., 2002 Actin cytoskeleton regulation in neuronal morphogenesis and structural plasticity. Annu Rev Cell Dev Biol 18: 601–635.

McMullan, R., and S. J. Nurrish, 2011 The RHO-1 RhoGTPase modulates fertility and multiple behaviors in adult C. elegans. PLoS One 6: e17265.

Montell, D. J., 1999 The genetics of cell migration in Drosophila melanogaster and Caenorhabditis elegans development. Development 126: 3035–3046.

Moore, S. W., M. Tessier-Lavigne and T. E. Kennedy, 2007 Netrins and their receptors. Adv Exp Med Biol 621: 17–31.

Morita, K., K. Hirono and M. Han, 2005 The Caenorhabditis elegans ect-2 RhoGEF gene regulates cytokinesis and migration of epidermal P cells. EMBO Rep 6: 1163–1168.

Motegi, F., and A. Sugimoto, 2006 Sequential functioning of the ECT-2 RhoGEF, RHO-1 and CDC-42 establishes cell polarity in Caenorhabditis elegans embryos. Nat Cell Biol 8: 978–985.

Nichol, R. I., K. M. Hagen, D. C. Lumbard, E. W. Dent and T. M. Gomez, 2016 Guidance of Axons by Local Coupling of Retrograde Flow to Point Contact Adhesions. J Neurosci 36: 2267–2282.

Norris, A. D., J. O. Dyer and E. A. Lundquist, 2009 The Arp2/3 complex, UNC-115/abLIM, and UNC-34/Enabled regulate axon guidance and growth cone filopodia formation in Caenorhabditis elegans. Neural Dev 4: 38.

Norris, A. D., and E. A. Lundquist, 2011 UNC-6/netrin and its receptors UNC-5 and UNC-40/DCC modulate growth cone protrusion in vivo in C. elegans. Development 138: 4433–4442.

Norris, A. D., L. Sundararajan, D. E. Morgan, Z. J. Roberts and E. A. Lundquist, 2014 The UNC-6/Netrin receptors UNC-40/DCC and UNC-5 inhibit growth cone filopodial protrusion via UNC-73/Trio, Rac-like GTPases and UNC-33/CRMP. Development 141: 4395–4405.

Patel, F. B., Y. Y. Bernadskaya, E. Chen, A. Jobanputra, Z. Pooladi et al., 2008 The WAVE/SCAR complex promotes polarized cell movements and actin enrichment in epithelia during C. elegans embryogenesis. Dev Biol 324: 297–309.

Petratos, S., Q. X. Li, A. J. George, X. Hou, M. L. Kerr et al., 2008 The beta-amyloid protein of Alzheimer’s disease increases neuronal CRMP-2 phosphorylation by a Rho-GTP mechanism. Brain 131: 90–108.

Rogers, S. L., U. Wiedemann, U. Hacker, C. Turck and R. D. Vale, 2004 Drosophila RhoGEF2 associates with microtubule plus ends in an EB1-dependent manner. Curr Biol 14: 1827–1833.

Rosslenbroich, V., L. Dai, S. L. Baader, A. A. Noegel, V. Gieselmann et al., 2005 Collapsin response mediator protein-4 regulates F-actin bundling. Exp Cell Res 310: 434–444.

Shakir, M. A., K. Jiang, E. C. Struckhoff, R. S. Demarco, F. B. Patel et al., 2008 The Arp2/3 activators WAVE and WASP have distinct genetic interactions with Rac GTPases in Caenorhabditis elegans axon guidance. Genetics 179: 1957–1971.

Shekarabi, M., and T. E. Kennedy, 2002 The netrin-1 receptor DCC promotes filopodia formation and cell spreading by activating Cdc42 and Rac1. Mol Cell Neurosci 19: 1–17.

Spencer, A. G., S. Orita, C. J. Malone and M. Han, 2001 A RHO GTPase-mediated pathway is required during P cell migration in Caenorhabditis elegans. Proc Natl Acad Sci U S A 98: 13132–13137.

Srayko, M., A. Kaya, J. Stamford and A. A. Hyman, 2005 Identification and characterization of factors required for microtubule growth and nucleation in the early C. elegans embryo. Dev Cell 9: 223–236.

Struckhoff, E. C., and E. A. Lundquist, 2003 The actin-binding protein UNC-115 is an effector of Rac signaling during axon pathfinding in C. elegans. Development 130: 693–704.

Sundararajan, L., M. L. Norris, S. Schoneich, B. D. Ackley and E. A. Lundquist, 2014 The fat-like cadherin CDH-4 acts cell-non-autonomously in anterior-posterior neuroblast migration. Dev Biol 392: 141–152.

Tessier-Lavigne, M., and C. S. Goodman, 1996 The molecular biology of axon guidance. Science 274: 1123–1133.

Van Goor, D., C. Hyland, A. W. Schaefer and P. Forscher, 2012 The role of actin turnover in retrograde actin network flow in neuronal growth cones. PLoS ONE 7: e30959.

Varadarajan, S. G., and S. J. Butler, 2017 Netrin1 establishes multiple boundaries for axon growth in the developing spinal cord. Dev Biol 430: 177–187.

Varadarajan, S. G., J. H. Kong, K. D. Phan, T. J. Kao, S. C. Panaitof et al., 2017 Netrin1 Produced by Neural Progenitors, Not Floor Plate Cells, Is Required for Axon Guidance in the Spinal Cord. Neuron 94: 790–799 e793.

Wadsworth, W. G., H. Bhatt and E. M. Hedgecock, 1996 Neuroglia and pioneer neurons express UNC-6 to provide global and local netrin cues for guiding migrations in C. elegans. Neuron 16: 35–46.

Wittmann, T., and C. M. Waterman-Storer, 2001 Cell motility: can Rho GTPases and microtubules point the way? J Cell Sci 114: 3795–3803.

Yamauchi, K., M. Yamazaki, M. Abe, K. Sakimura, H. Lickert et al., 2017 Netrin-1 Derived from the Ventricular Zone, but not the Floor Plate, Directs Hindbrain Commissural Axons to the Ventral Midline. Sci Rep 7: 11992.

Yan, J., D. L. Chao, S. Toba, K. Koyasako, T. Yasunaga et al., 2013 Kinesin-1 regulates dendrite microtubule polarity in Caenorhabditis elegans. Elife 2: e00133.

Yang, Y., W. S. Lee, X. Tang and W. G. Wadsworth, 2014 Extracellular matrix regulates UNC-6 (netrin) axon guidance by controlling the direction of intracellular UNC-40 (DCC) outgrowth activity. PLoS One 9: e97258.

Yau, D. M., N. Yokoyama, Y. Goshima, Z. K. Siddiqui, S. S. Siddiqui et al., 2003 Identification and molecular characterization of the G alpha12-Rho guanine nucleotide exchange factor pathway in Caenorhabditis elegans. Proc Natl Acad Sci U S A 100: 14748–14753.

Zhang, X. F., A. W. Schaefer, D. T. Burnette, V. T. Schoonderwoert and P. Forscher, 2003 Rho-dependent contractile responses in the neuronal growth cone are independent of classical peripheral retrograde actin flow. Neuron 40: 931–944.

